# The Antimicrobial Efficacy Of Plasma Activated Water Is Modulated By Reactor Design And Water Composition

**DOI:** 10.1101/2021.07.14.452435

**Authors:** Joanna G. Rothwell, David Alam, Dee A. Carter, Behdad Soltani, Robyn McConchie, Renwu Zhou, Patrick J. Cullen, Anne Mai-Prochnow

## Abstract

Plasma activated water (PAW) contains a cocktail of reactive oxidative species and free radicals and has demonstrated efficacy as a sanitizer for fresh produce, however there is a need for further optimization. The antimicrobial efficacy of PAW produced by a bubble spark discharge (BSD) reactor and a dielectric barrier discharge-diffuser (DBDD) reactor operating at atmospheric conditions with air, discharge frequencies of 500, 1000 and 1500 Hz, and MilliQ and tap water, was investigated with model organisms *Listeria innocua* and *Escherichia coli*. Optimal conditions were subsequently employed for pathogenic bacteria *Listeria monocytogenes, E. coli* and *Salmonella enterica*. PAW generated with the DBDD reactor reduced more than 6-log CFU of bacteria within 1 minute of treatment. The BSD-PAW, while attaining high CFU reduction was less effective, particularly for *L. innocua*. Analysis of physicochemical properties revealed BSD-PAW had a greater variety of reactive species than DBDD-PAW. Scavenger assays were employed to specifically sequester reactive species, including the short-lived superoxide (·O_2_^-^) radical that could not be directly measured in the PAW. This demonstrated a critical role of superoxide for the inactivation of both *E. coli* and *L. innocua* by DBDD-PAW, while in BSD-PAW it had a role in *L. innocua* inactivation only. Overall, this study demonstrates the potential of DBDD-PAW in fresh produce, where there is a need for sterilization while minimizing chemical inputs and residues and maintaining food quality. Highly effective PAW was generated using air as a processing gas and tap water, making this a feasible and cost-effective option.

**Importance:** There is a growing demand for fresh food produced with minimal processing, however guaranteeing microbial safety in the absence of a thermal kill step is challenging. Plasma-activated water (PAW) is a promising novel antimicrobial but its use in high-risk applications like the sanitization of fresh produce requires further optimization. This study demonstrated the importance of reactor design in the production of reactive species in PAW with capacity to kill bacteria. Very effective PAW was generated using a dielectric barrier discharge-diffuser (DBDD) system, with antimicrobial activity attributed to the presence of superoxide radicals. The DBBD reactor used air as a processing gas and tap water, highlighting the potential of this approach as a cost-effective and green alternative to chemical treatment methods that are currently used in food decontamination.

## Introduction

Developing safe and effective antimicrobial technologies is of great importance in many areas where unwanted microbial growth can occur, including medical, water and food industries. Plasma activated water (PAW) is a promising antimicrobial tool with a wide variety of potential applications that reduce the need for toxic chemicals. Of particular interest is the application of PAW in fresh produce processing, since current disinfection methods rely on chemicals that have significant health and environmental impacts. PAW has been shown to reduce pathogenic microbes with relevance to fresh produce safety, including *Escherichia coli* (1, 2), *Staphylococcus aureus* (3–7), *Bacillus. subtilis* (8), *Listeria monocytogenes* and *Salmonella Typhimurium* (9), while retaining product quality (10).

PAW can be produced from cold plasma, a non-thermal plasma composed of highly energetic particles including UV, charged ions, accelerated electrons, radical species, excited molecules and atoms (11, 12). Cold plasma discharged over the surface of water or directly ignited within water results in plasma-liquid interactions that generate various reactive species with antibacterial activity (13). These include reactive oxygen species, such as hydroxyl radicals (•OH), atomic oxygen, superoxide (O_2_^-^), ozone (O_3_) and hydrogen peroxide (H_2_O_2_), as well as reactive nitrogen species including atomic nitrogen, peroxynitrite, nitric oxide, nitrates (NO_3_^-^) and nitrites (NO_2_^-^) (14). While most reactive species are extremely short-lived, nitrite, nitrate and hydrogen peroxide are relatively stable and can last several months depending on storage conditions; for example they can be preserved by freezing (6, 11). PAW has an acidic pH due to hydrogen peroxide, nitric and peroxynitrous acid, and a high oxidation-reduction potential (ORP) largely due to hydrogen peroxide (1, 15). The individual and synergistic effects of these physicochemical properties and the reactive species in this dynamic mixture, make it potently antimicrobial (16).

There are multiple PAW configurations that have been described in depth (13). To date, the most common configuration to produce PAW has been to discharge gaseous plasma across water surfaces (17), but this confers a relatively poor transfer of reactive species. More recent configurations use multiphase discharges where plasma gas is bubbled through water, allowing a substantially greater transfer of reactive species into the water. Two such methods are shown in Figure 1. The bubble spark discharge (BSD) transmits reactive species into the water via high-intensity spark plasma that creates reactive species at the plasma-water interface, in conjunction with plasma gas bubbles that diffuse reactive species into the water. In the dialectic barrier discharge diffusor (DBDD), the reactor produces a larger volume of lower energy DBD plasma that is introduced into the solution via a diffusor, producing copious small bubbles with a large surface area to allow mass transfer of reactive species into solution. Different discharge schemes can be employed to regulate the concentration and types of reactive species generated in PAW (18). However, further research is required to understand how to maximize the production of reactive species using the different plasma reactor designs and to determine how these work to inhibit and kill bacterial cells.

**Figure 1.**
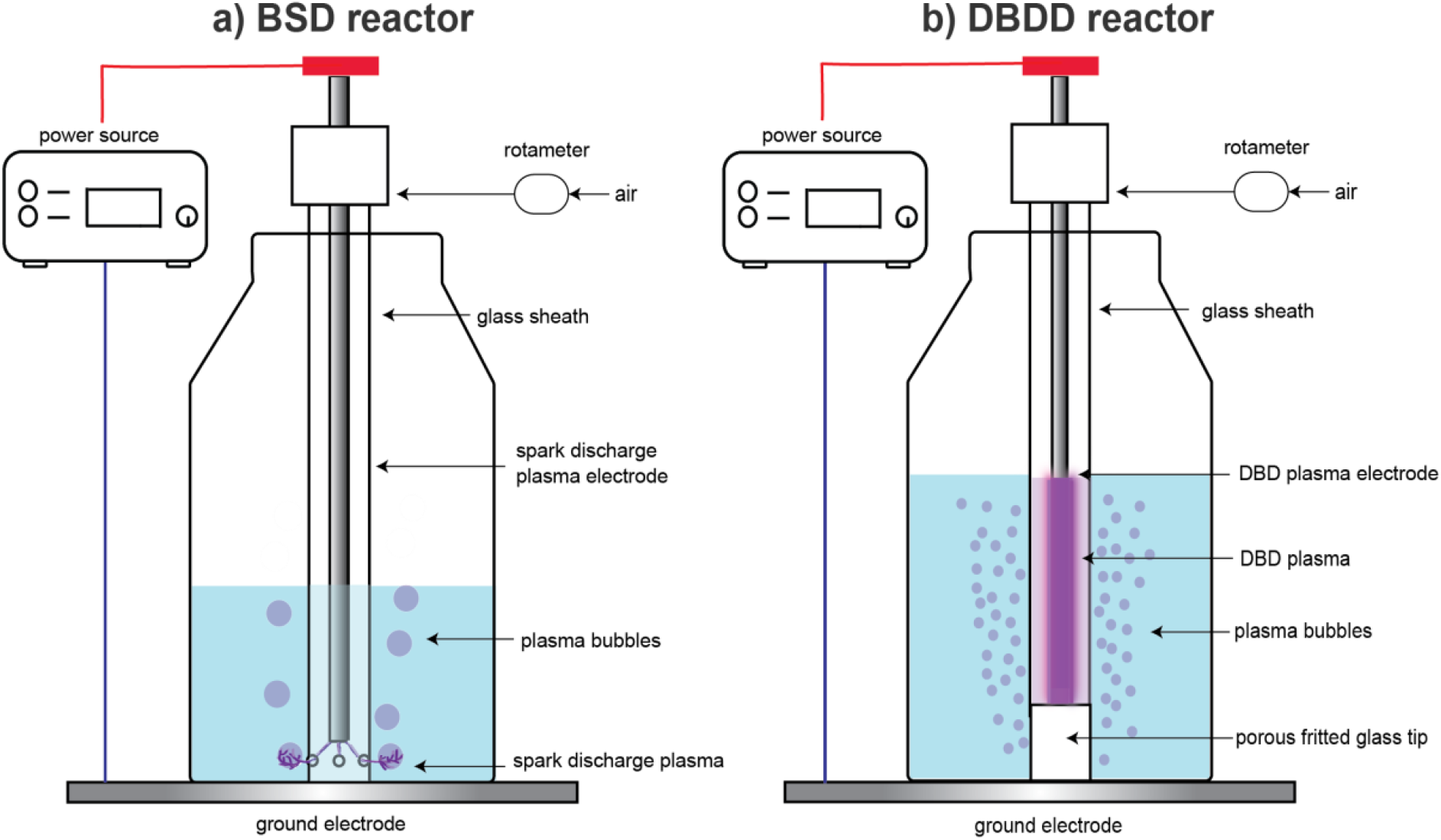
Schematic of the bubble spark discharge (BSD) reactor a) and the dielectric barrier discharge diffusor (DBDD) reactor b) for PAW production.

The current study set out to compare BSD and DBBD PAW systems for their antimicrobial efficacy against representative bacterial food pathogens. While most PAW research has used sterile distilled and deionized water (19), we also tested tap water as a more economically viable option for food processing. We then quantified the reactive species produced by the different systems and used selective quenching agents to determine which reactive species contributed most to the antimicrobial activity.

## 2.0 Materials and Methods

### 2.1 Plasma reactor design

BSD and DBDD reactors were designed in-house (Plasmaleap Technologies, Sydney, Australia) using the setup shown in Figure 1. For both reactors, power was supplied by a high voltage micropulse generator Leap100 (PlasmaLeap Technologies, Sydney, Australia), capable of providing a high voltage pulse of up to 80 kV (p-p), a repetitive pulse frequency from 100 to 3000 Hz, and maximum output power of 400 W. Both reactors used stainless steel high voltage electrodes fitted in place by machined acrylic spacers and polytetrafluoroethylene (PTFE) tee fittings. The PTFE tee fitting connected to the open end of the dispersion tube in both reactors, and was used to supply the airflow into the tubes as well as to support the electrode. A rotameter to control the airflow rate was attached to a machined Teflon fitting on the outside tubes. The high voltage electrode from the high voltage supply was connected directly to the stainless steel electrode, whereas the ground electrode consisted of a stainless steel plate under the bottle connected to the ground wire from the power supply. Both reactors were placed into 250 mL Schott bottles filled with autoclaved tap or MilliQ water.

The body of the BSD reactor consisted of a 175 mm-length quartz tube (10 mm outside diameter (OD) and 1.5 mm wall thickness) with one end sealed. Four 0.4 mm holes were positioned radially approximately 5 mm above the sealed end. The electrode was a stainless steel rod (4 mm OD) inserted coaxially along the length of the quartz tube. The DBDD reactor employed a conventionally coaxial electrode configuration connected to a glass diffusor to transfer the reactive gas species generated in the plasma into the water. The body of the DBDD reactor consisted of a 150 mm long gas dispersion tube with a fritted glass tip with a pore size 10-20 μm (Ace Glass 720208). A sealed borosilicate tube (≈4mm OD, 0.8mm wall thickness) was used as a dielectric insulator to sheath the high voltage electrode inside the length of the dispersion tube.

### 2.2 Bacterial cultures

The bacterial species used in this study are listed in Table 1. Model non-pathogenic surrogate species *Listeria innocua* (ICMPR:80-16-328-502) and *E. coli* (ATCC R25922) were used to observe the antimicrobial activity of PAW under different plasma conditions. These conditions were then applied to isolates of pathogenic bacteria linked to foodborne illness cases, which were sourced from the New South Wales Enteric Reference Laboratory at the Centre for Infectious Diseases and Microbiology Laboratory Services (CIDMLS), NSW Health Pathology, Westmead. Bacteria were resuscitated from glycerol stocks stored at −80 °C using tryptic soy agar (TSA: 17 g L^−1^ pancreatic digest of casein, 5 g L^−1^ Papaic digest of soybean meal, 5 g L^−1^ sodium chloride, 15 g L^−1^ agar-agar) plates for *Salmonella* spp. and *E. coli* and tryptic soy sheep blood agar (TSBA: TSA with 5 % defibrinated sheep’s blood) for *Listeria* spp., and incubated at 37 °C overnight. A single colony was then inoculated into 10 mL of tryptic soy broth (TSB, 17 g L^−1^ pancreatic digest of casein, 2.5 g L^−1^ D(+)glucose monohydrate, 3 g L^−1^ Papaic digest of soybean meal, 5 g L^−1^ sodium chloride, 2.5 g L^−1^ di-potassium hydrogen phosphate) for 24 hours with shaking at 30 °C for *Listeria* spp. and 37 °C for *Salmonella* spp and *E. coli*. 10 μL from each suspension was then transferred into 10 mL of TSB and incubated under the same conditions for 18 hours. Cultures were harvested by centrifugation at 3000 rpm for 10 minutes at 4 °C and the cell pellet was resuspended in a volume of PBS to obtain a concentration of ~1 × 10^8^ CFU mL^−1^. The final inoculum was stored at 4 °C and used within 4 hours. The final concentration was verified by back-plating, where *Listeria* solutions were spread onto TSBA and incubated for 48 h at 30 °C, and *E. coli* was spread onto TSA and incubated for 24 h at 37 °C.

**Table 1.** Bacterial isolates subjected to PAW treatments

| Bacterial Species | Strain designation | Serotype | Source (Year) |
| --- | --- | --- | --- |
| *Listeria innocua* | ICMPR:80-16-328-5025 (non pathogenic) | - | Environmental (2017) |
| *Listeria monocytogenes* | ICPMR: 07-15-147-1471 | 4b | Ascetic Fluid (2015) |
|  | ICMPR: 80-13-220-4103 | 1/2b | Blood (2013) |
| *Escherichia coli* | ATCC R25922 (non pathogenic) | (Migula) Castellani and Chalmers 1919 | Human clinical isolate (1977) |
|  | ICMPR: 40-16-302-2227 | O157:H7 | Faeces (2016) |
|  | ICMPR: 80-16-270-5374 | O26: H11 | Faeces (2016) |
| *Salmonella enterica* subsp. *Enterica* | ICMPR: 06-17-184-1802 | Saintpaul | Faeces (2017) |
|  | ICMPR: 80-17-173-5603 | Hvittingfoss | Faeces (2017) |

### 2.3 Bacterial Inhibition by PAW

The inhibition assay was first optimized using non-pathogenic bacteria. For both reactors we tested three different discharge frequencies (500, 1000 and 1500 Hz) using either autoclaved tap water or Milli-Q water. Air provided to the reactors at 1 standard liter per minute (SLM) and plasma treatments were performed within a fume hood. For the BSD reactor, a 100 μL aliquot of each inoculum prepared as above was pipetted into 100 mL of MilliQ or tap water in the 250 mL Schott bottle (figure 1). The setting used on the power supply for the BSD were 150 V, 100 μs duty cycle 60 kHz resonance frequency. For the DBDD reactor, 200 μL of the inoculum was pipetted into 200 mL of Milli-Q or tap water in the 250 mL Schott bottle. The power supply settings used were the same as those used with the BSD reactor except voltage, which was 120 V. Different volumes of water were used in the two reactor systems, as the BSD reactor generates a higher intensity plasma and requires a low volume of water to concentrate the reactive species, while the DBDD reactor requires a larger volume to provide a greater surface area for the bubbles to exchange reactive species with the water. Both reactors were run for 12.5 minutes with samples removed at 1, 2.5, 5 7.5, 10 and 12.5 minutes. A control was included for both reactors where the same volume of water and inoculum were bubbled under identical airflow conditions for 12.5 minutes without the power source turned on. Each treatment was neutralized after plasma exposure by dilution into a modified Dey-Engley neutralizing broth (5 g L^−1^ pancreatic digest of casein, 2.5 g L^−1^ yeast extract, 10 g L^−1^ D(+)glucose monohydrate, 5 g L^−1^ polysorbate 80, 7 g L^−1^ lecithin, 6 g L^−1^ sodium thiosulfate, 2.5g L^−1^ sodium bisulfite). Surviving bacteria were serially diluted in a 96-well plate with 0.1 % peptone water and enumerated using spread plates as above. All experiments were performed in duplicate, with three biological replicates performed on three separate days.

Once the optimal plasma generating conditions had been determined, the six strains of pathogenic bacteria (Table 1) were tested using the DBDD reactor with tap water at 1000 Hz and 120 V for 1 and 2 minutes.

### 2.4 Measurement of PAW Composition

Concentrations of ozone (O_3_), hydrogen peroxide (H_2_O_2_), nitrite (NO_2_^-^) and nitrate (NO_3_^-^) along with pH, temperature and conductivity were assessed in the PAW produced by the BSD and DBDD reactors over 12.5 min. Ozone was measured using a Hanna Instrument multiparameter photometer (H183399, Rhode Island USA) with colorimetric ozone kits. H_2_O_2_ was measured by a titanium sulfate method (20). Nitrite (NO_2_^-^) was assessed using the Griess Reagent method (21), and nitrate was quantified by High-Pressure Liquid Chromatography (HPLC) using a Shimadzu Prominence-i HPLC system equipped with a photodiode array detector. A YMC-Pack C18 column (4.6 mm × 250 mm, 5 μm) operated at 40 °C was used with a mobile phase of 10:90 of methanol/0.1 % acetic acid (22) in Milli-Q water at a flow rate of 0.5 mL min^−1^ for 10 min with a 10 μL injection volume. The concentration of nitrate was measured at an absorbance of 200 nm, with a calibration curve plotted by measuring absorbance of a series of standard solutions at 1, 5, 10, 25, 30, 40, 50, 60, 100 mg/L. PAW samples were initially passed through a filter syringe (13 mm diameter, 0.45 um pore size) before HPLC analysis. Temperature was measured using a Tinytag TGP-4200 Data Logger (Gemini, United Kingdom) with corresponding SWCD-0040: Tinytag Explorer software. pH was measured using a SevenCompact S220 pH meter (Mettler-Toldeo, Switzerland). Conductivity of PAW was measured using a four-ring conductivity probe (HI76312) connected to the Hanna Instrument multiparameter photometer (H183399, Rhode Island, USA). Optical emission spectra (OES) measurements were recorded for both reactors using an Andor Shamrock SR-500i-A-R spectrometer. This emission spectroscopy technique determines how much of an individual excited atom or ion is present in a plasma via the characteristic wavelengths of electromagnetic radiation emitted by individual species. Typical spectra emitted from different reactors were used to investigate the main excited active species generated by air plasma ranging from 200 to 900 nm. The optical fiber was located at the DB section in the DBDD reactor or at the same level as the bubble-liquid interface in the BSD reactor. Chemical analysis of the tap water was performed by Envirolab Sevices (Sydney, Australia), where the following water properties were assessed: pH, electrical conductivity, total dissolved solids, ionic balance (calcium, potassium, magnesium, sodium, hardness, hydroxide alkalinity, bicarbonate alkalinity, carbonate alkalinity, total alkalinity, chloride, sulfate), fluoride, nitrate, nitrite, total metals (arsenic, cadmium, copper, lead, manganese, iron) and dissolved iron.

### 2.5 Scavenger Assay

Chemical scavengers were used to specifically quench reactive species generated in the PAW to investigate their roles in antimicrobial activity. Scavenger types and concentrations were selected based on previous studies in PAW and included 20 μM hemoglobin for nitric oxide (NO) (23), 200 mM mannitol for hydroxide (·OH) (24), 10 mM sodium pyruvate for hydrogen peroxide (H_2_O_2_) (21) 100 uM Uric acid for ozone (O_3_) (21) and 20 mM tiron for superoxide ion (·O_2_^-^) (24) (Sigma-Aldrich, Missouri, United States).

Scavenger solutions and bacterial suspensions at a final concentration of ~1 × 10^5 cells / mL (section 2.1) were added to the water in the 250 mL Schott bottle prior to plasma activation. Reactor conditions were selected for the scavenger assay based on the results of antimicrobial activity analysis, such that there was either a complete reduction of bacteria, or if this was not achieved the longest run time with the highest discharge frequency was used. For the BSD reactor, for *L. innocua* this was 1500 Hz for 5 min in milli-Q or for 13.5 min in tap water, and for *E. coli* it was 1500 Hz for 2 min in milli-Q or for 12.5 min in tap water. For the DBDD reactor, 1000 Hz for 1 min was used for both water types and both bacteria. After plasma treatment, aliquots of the solutions were removed and immediately neutralized in DE broth before being serially diluted and plated out as described above (section 2.1).

### 2.6 Statistical analysis

Two-way ANOVAs followed by Dunnett’s multiple comparisons tests comparing the PAW treatments to the water controls (Fig. 2) or comparing scavenger treatments to the no scavenger PAW condition (Fig. 6) were performed using GraphPad Prism version 8.0.0 (GraphPad Software, San Diego, California USA). A p-value of <0.05 was considered statistically significant.

**Figure 2:**
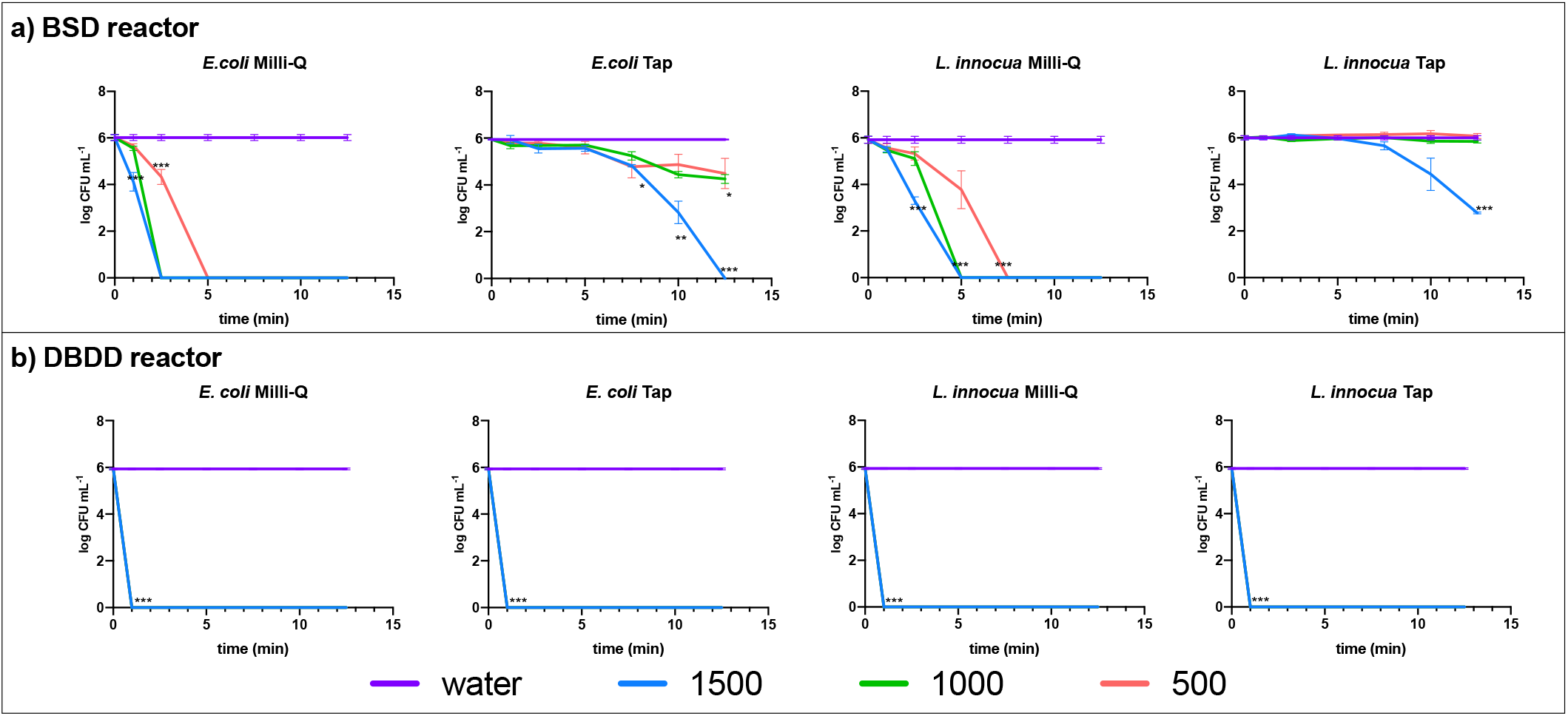
Reduction of bacterial CFU using the BSD reactor a) or the DBDD reactor b) with three discharge frequencies over 12.5 minutes. A p-value of <0.05 was considered statistically significant.

## 3.0 Results

### 3.1 PAW generated using BSD or DBDD reactors with Milli-Q or tap water is antibacterial with variable efficacy

The two different plasma generating probes (Fig. 1) were compared for their ability to reduce CFU numbers of *E. coli* and *L. innocua* using either milliQ or tap water (Fig. 2). Chemical analysis of tap water indicated a variety of different inorganic ions were present, including trace amounts of various metals (supplementary table 1). PAW generated by the DBDD reactor (PAW-DBDD) was the most effective, achieving a complete (6-log) reduction of both bacterial species within 1 minute with both tap and MilliQ water and across all discharge frequencies. PAW produced by the BSD reactor (BSD-PAW) did not reduce the bacterial load as rapidly as the DBDD-PAW under any of the conditions tested, and more variability was observed across treatments. In all cases, the 1500 Hz discharge frequency was the most effective, and CFU were reduced more rapidly using MilliQ than tap water. Treatment response also varied by bacterial species: BSD-PAW generated using MilliQ water reduced *E. coli* counts by 6 log in 2.5 minutes at 1000 or 1500 Hz, and in 5 minutes at 500 Hz, while it took an additional 2.5 minutes to achieve the same reduction of *L. innocua*. For the BSD-PAW generated using tap water, *E. coli* was reduced by 6 log after 12.5 minutes of treatment at 1500 Hz, but the highest reduction for *L. innocua* was only 3 log.

As DBDD-PAW was clearly more efficacious, six foodborne pathogenic bacteria that have caused severe disease in humans (Table 1) were tested using this system at 1000 Hz, 120 V with tap water. All six bacterial strains were completely inactivated (6 log) within 1 minute (Fig. 3).

**Figure 3:**
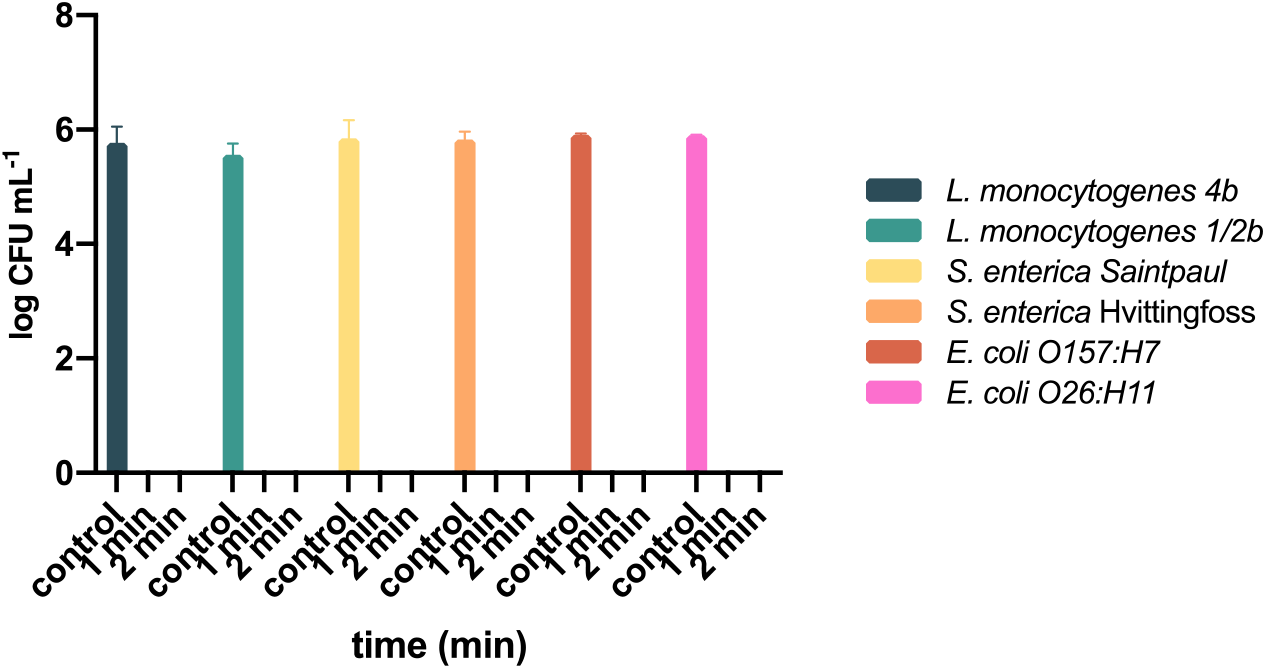
Complete reduction of pathogenic bacterial species is achieved using the DBDD reactor with conditions optimized for non-pathogenic strains (1000 Hz and 120 V with tap water).

### 3.2 Physicochemical properties of PAW varied with reactor design, generation conditions, and water type

The physicochemical properties of the plasma and the PAW were assessed for each reactor. Optical emission spectra (OES) were used to determine the main excited active species generated by each type of plasma in the air. As these initial short-lived species are highly reactive and cause downstream reactions that result in the generation of other secondary species, the characterization of OES is critical for defining the physicochemical properties of the PAW. The OES obtained from the DBDD, and BSD reactors ranged from 200 to 900 nm and had large nitrogen emission peaks, indicating that high levels of excited nitrogen molecules and ions were present (Fig. 4). Excited oxygen emission peaks were observed for both reactors, with slightly higher peaks for the BSD reactor. Hydroxy (·OH) emission peaks were only observed in the plasma produced by the BSD reactor.

**Figure 4.**
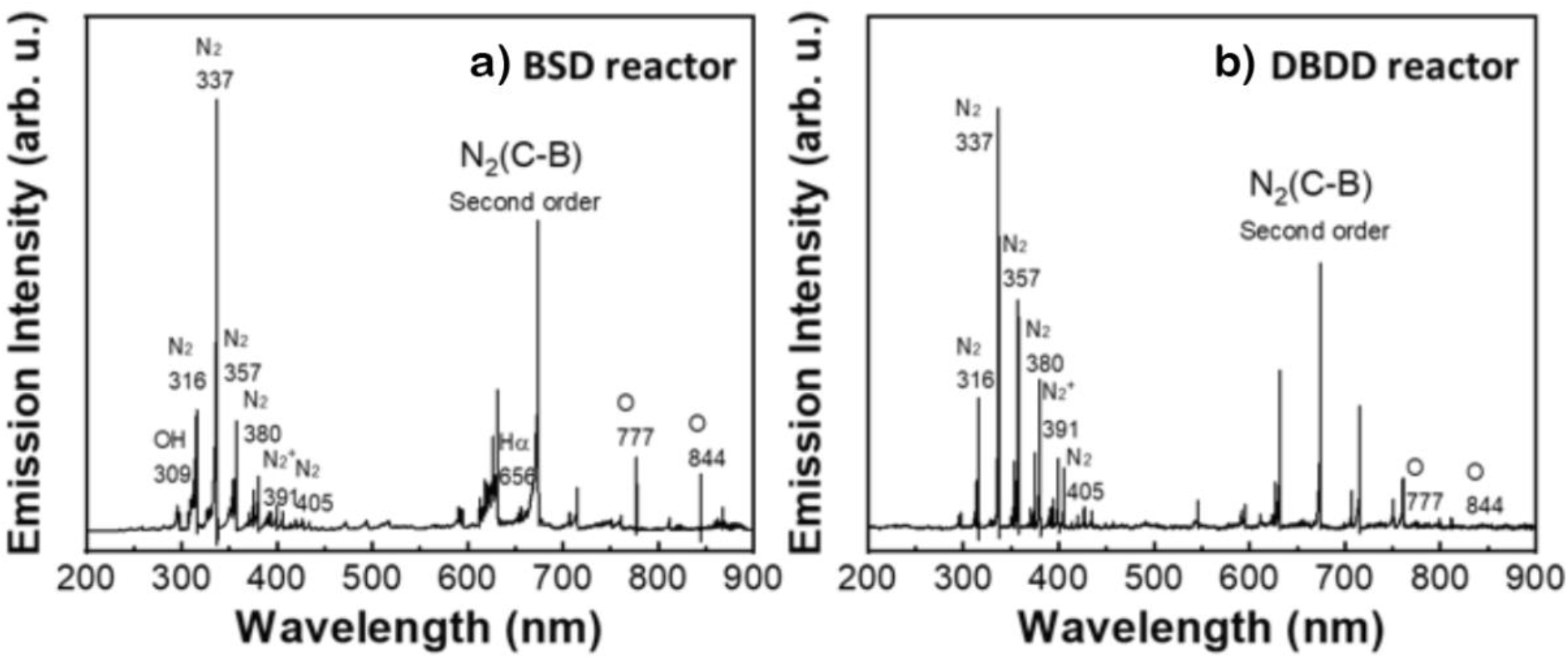
Optical emission spectra of the a) BSD and b) DBDD reactors measured using optical fiber indicated the presence of reactive nitrogen and oxygen species in the plasma that they generated.

Reactive species produced in PAW over time were quantified using the same parameters that had been applied in the tests for antibacterial activity (Fig. 5). PAW generated by the BSD reactor contained higher concentrations of all reactive species tested than DBDD-PAW. In BSD-PAW, all measured reactive species increased over time, while in DBDD-PAW, only NO_3_^-^ and small amounts of O_3_ were recorded. For O_3_, the BSD-PAW had maximum measurable concentrations of ~2 mg/L and levels increased with the reactor discharge frequency, while the DBDD-PAW had substantially lower amounts of O_3_ and the lowest discharge frequency produced the highest O_3_ levels with both tap and Milli-Q water. H_2_O_2_ and NO_2_^-^ were not present in DBDD-PAW, while in the BSD-PAW these reactive species increased linearly with BSD activation time. NO_3_^-^ increased linearly over time in PAW generated by both reactors, with the highest discharge frequency producing the highest levels. The physicochemical properties of the PAW produced by the two reactors also differed. The pH of BSD-PAW was lower than DBDD-PAW, while the temperature and conductivity of BSD-PAW were higher. Overall, the physicochemical properties of BSD-PAW appeared more extreme than the DBDD-PAW. The water used in the generation of PAW strongly affected its properties. pH decreased rapidly when Milli-Q water was used, while it remained stable with the use of tap water (Fig. 5). Temperature was not affected by water type, however it became substantially higher in the DBS reactor and increased with discharge frequency. Conductivity was also more affected in MilliQ than tap water, particularly with the DBS reactor.

**Figure 5.**
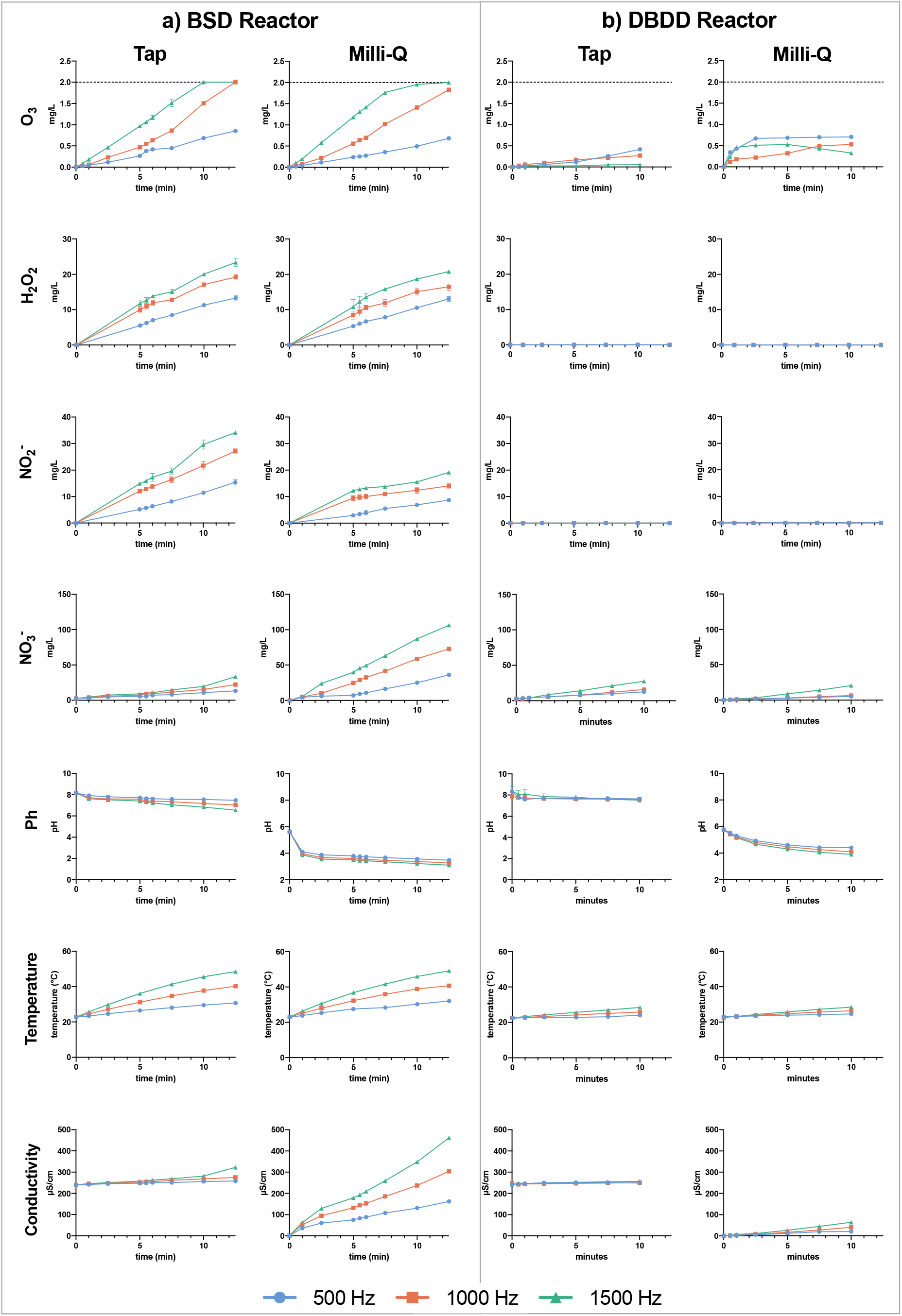
The concentration of reactive species present in PAW produced by the BSD reactor a) and DBDD reactor b) using either tap or milli-Q water and run over a 12.5 minute time course. Higher concentrations and a more complex mix of reactive species was observed in the BSD-PAW. Reactive species generally increased linearly over time, with higher discharge frequencies mostly leading to higher concentrations of reactive species in the PAW generated by both reactors.

### 3.3 The scavenging of superoxide led to survival of bacteria in DBDD-PAW

To test the role of some of the individual reactive species present in PAW, including some that could not be directly measured, we used a range of scavengers to selectively remove particular species, including tiron (which scavenges superoxide), mannitol (hydroxyl radical), uric acid (ozone), hemoglobin (nitric oxide) and sodium pyruvate (hydrogen peroxide). The addition of tiron led to significantly increased survival of *L. innocua* in both the BSD-PAW and the DBDD-PAW, and increased *E. coli* survival in DBDD-PAW. There was no significant difference in the scavenging results for the PAW generated in Milli-Q or tap water for both reactors. None of the other scavengers affected the survival of the bacterial pathogens.

## 4.0 Discussion

This study compared the efficacy of PAW generated by two novel plasma reactors against foodborne organisms. PAW generated using the DBDD reactor clearly displayed more rapid and acute antimicrobial activity than the BSD-PAW. The DBDD-PAW led to a ~6-log reduction of all pathogenic and non-pathogenic bacteria tested within 1 minute, and this occurred with both tap and Milli-Q water (Fig. 2). In contrast, using the BSD-PAW, a 6-log reduction of *E. coli* and *L. innocua* in Milli-Q water took 2.5 and 5 minutes, respectively. Tsoukou *et. al*. (25), who employed a system where the plasma was generated 5 mm above deionized water, also reported greater bactericidal activity of PAW generated using a dialectic barrier discharge system compared to a spark discharge system.

Analysis of the reactive species produced in PAW found these were very different between the two reactors, with fewer types and lower levels of measured reactive species in the DBDD-PAW despite its greater antimicrobial power (Fig. 5). However, the addition of tiron completely quenched the antibacterial activity of the DBDD-PAW (Fig. 6), indicating that superoxide ions and/or downstream reactions from this reactive species were primarily responsible for its powerful antimicrobial activity. Trion selectively interacts with and is oxidized by superoxide, allowing its effective removal from the solution (26, 27). Our finding is supported by previous research showing that superoxide anions and downstream reactive species such as singlet molecular oxygen are significant contributors to the antimicrobial activity of PAW produced by dialectic barrier discharge against yeast cells (28) and *Salmonella Typhimurium* (29).

**Figure 6.**
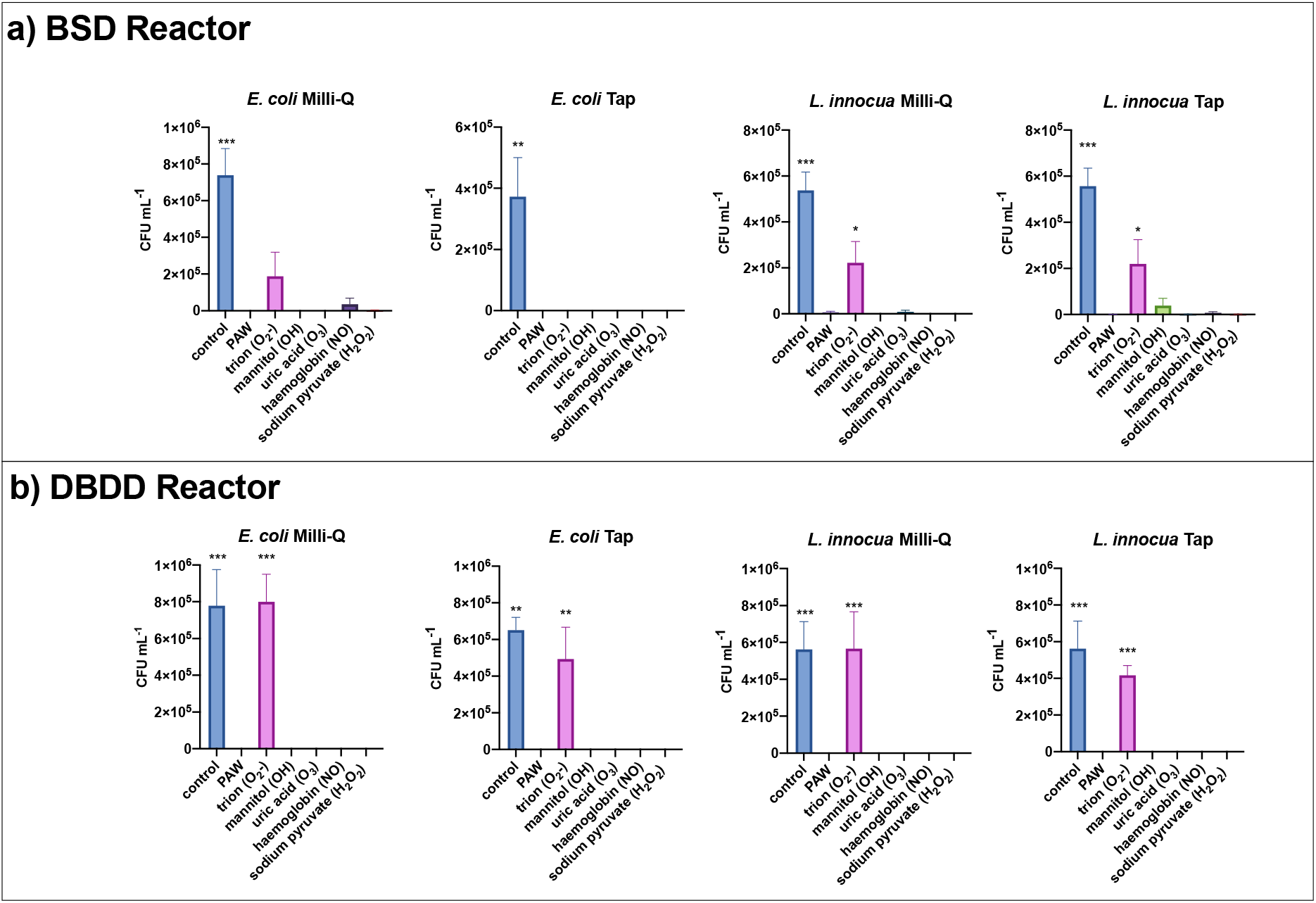
Addition of reactive species scavengers altered the antimicrobial efficacy of BSD-PAW a) and DBDD-PAW b). The addition of the superoxide scavenger trion led to increased survival across most of the treatments, but for BSD this only reached statistical significance with *L. innocua* treatment. None of the other scavengers led to increased bacterial survival.

When PAW was made with the BSD reactor, the addition of tiron lead to increased but not complete survival of *L. innocua*, indicating that superoxide contributes to the ability of BSD-PAW to kill *Listeria*, but it is not as critical as in DBDD-PAW. With BSD-PAW there were no distinct scavengers that sequestered antimicrobial activity against *E. coli*. Other reactive species that could not be elucidated in this study, or synergy between components of the complex mixture present in BSD-PAW, may have been responsible for its bactericidal activity. In addition, recent studies suggest that plasma treatment elicits physical conditions, including electric fields (30), that were not investigated here. Further research is required to fully understand what causes the biocidal effects of BSD-PAW.

The analysis of reactive species present in the PAW highlighted the diversity of molecular species that can be produced by plasma, particularly when using the spark discharge system. The OES (Fig. 4) varied between the two systems due to the different excited nitrogen and oxygen species generated in the gaseous plasmas, and these correspond to the different profiles of reactive species seen in the PAW (Fig. 5). For example, H_2_O_2_ was only quantified in BSD-PAW (Fig. 5), and this can be attributed to the generation of ·OH radicals (Fig. 4) produced directly at the plasma-water interface of the BSD that diffuse into solution and react to form hydrogen peroxide (31).. Similarly, a higher concentration of NO_2_^-^ and NO_3_^-^ were detected in the BSD-PAW compared to the DBDD-PAW (Fig. 5) as the BSD reactor generated a higher intensity plasma which is able to form NOx gases (Fig. 4) that diffuse into the water to produce these reactive nitrogen species in solution. ROS concentrations were also higher in the BSD-PAW. The energetic collisions of electrons with O_2_ molecules (O_2_ + e^-^ → 2O + e^-^) account for the formation of ·O radicals (13) at 777.2 and 844.6 nm, which were produced by both reactors but in higher concentrations in the BSD reactor, resulting in higher ROS concentrations in the PAW generated by this reactor (Fig. 5).

Our findings have importance for translation in the fresh produce industry. Fresh produce treatment requires high water volumes, however most studies to date on PAW have used distilled or deionized water (32), and there is a paucity of research on the antimicrobial efficacy of tap PAW. Deionising water is not economically feasible for use in sanitation processes, and our study found Milli-Q water rapidly acidified to a pH of ~3-4 (Fig. 5) which may damage fresh produce. Water hardness has been shown to affect the active components of PAW, however (33), and as the water used in this study is relatively soft (supplementary data table 1), further work is required to determine whether the antimicrobial activity demonstrated here extends to DBDD-PAW produced using tap water of different quality.

The BSD-PAW, although less effective than DBDD-PAW in the current study, could still have applications in the food industry. The reactive species produced by this reactor were longer-lived compared to the DBDD-PAW, which largely relied on the short-lived reactive species superoxide, and BSD-PAW may be more suitable for use where there is a delay between PAW generation and use. The reactive species in the BSD-PAW increased linearly over time, and longer activation and treatment times may enhance its antimicrobial power. This could also be further increased by synergizing with existing antimicrobials such as hydrogen peroxide (34) or using different processing gas sources such as oxygen (35).

The high efficacy of DBDD-PAW against a variety of pathogenic bacterial strains when made with tap water and air as a processing gas, rather than more expensive gases such as argon or oxygen, indicates the potential of this technology as an economically viable *in situ* sanitation process. Moving to apply this to produce will introduce new challenges, however, as organic load due to soil or plant exudates may interfere with PAW activity. Determining whether these findings extend to additional pathogenic and spoilage species, including yeasts and molds, is also required. This study lays for foundation for further testing and development of PAW as a cost-effective and non-toxic alternative to methods that are currently used in food decontamination.

## Conclusions

PAW is a rapidly emerging technology with many promising applications. This study involved a comparison of the efficacy of novel PAW bubbler reactors in their bactericidal performance *in situ* using both tap and Milli-Q water. The DBDD reactor produced the most effective PAW, which successfully decontaminated 6-log of pathogenic bacteria in tap water within 1 minute, and superoxide was determined to be a critical reactive species for antimicrobial activity. Different generation conditions and water types produced different reactive species profiles that varied in bactericidal power when using BSD-PAW. Tap water is used in many industrial processes, and the strong bactericidal activity of tap PAW produced by the DBDD reactor provides a valuable platform for further research. This study has provided new insight into the potential of PAW as a sanitizing agent against industrially relevant bacterial species.

## Supplementary data

### Tap water analysis

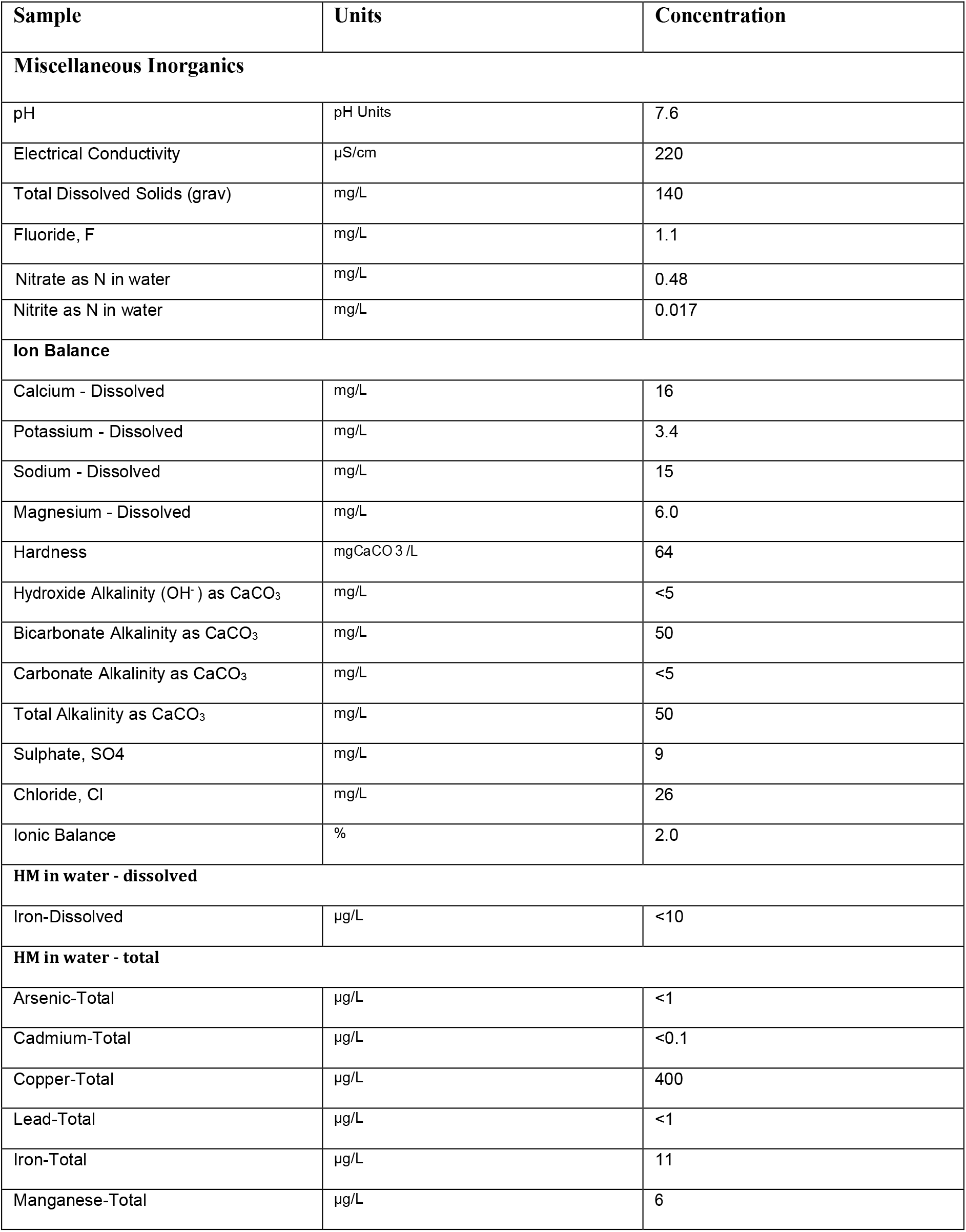

| Sample | Units | Concentration |
| --- | --- | --- |
| **Miscellaneous Inorganics** |  |  |
| pH | pH Units | 7.6 |
| Electrical Conductivity | µS/cm | 220 |
| Total Dissolved Solids (grav) | mg/L | 140 |
| Fluoride, F | mg/L | 1.1 |
| Nitrate as N in water | mg/L | 0.48 |
| Nitrite as N in water | mg/L | 0.017 |
| **Ion Balance** |  |  |
| Calcium - Dissolved | mg/L | 16 |
| Potassium - Dissolved | mg/L | 3.4 |
| Sodium - Dissolved | mg/L | 15 |
| Magnesium - Dissolved | mg/L | 6.0 |
| Hardness | mgCaCO<sub>3</sub>/L | 64 |
| Hydroxide Alkalinity (OH<sup>-</sup>) as CaCO<sub>3</sub> | mg/L | <5 |
| Bicarbonate Alkalinity as CaCO<sub>3</sub> | mg/L | 50 |
| Carbonate Alkalinity as CaCO<sub>3</sub> | mg/L | <5 |
| Total Alkalinity as CaCO<sub>3</sub> | mg/L | 50 |
| Sulphate, SO<sub>4</sub> | mg/L | 9 |
| Chloride, Cl | mg/L | 26 |
| Ionic Balance | % | 2.0 |
| **HM in water - dissolved** |  |  |
| Iron-Dissolved | µg/L | <10 |
| **HM in water - total** |  |  |
| Arsenic-Total | µg/L | <1 |
| Cadmium-Total | µg/L | <0.1 |
| Copper-Total | µg/L | 400 |
| Lead-Total | µg/L | <1 |
| Iron-Total | µg/L | 11 |
| Manganese-Total | µg/L | 6 |

